# Subcellular localization of microcystin in the liver and the gonad of medaka fish acutely exposed to microcystin-LR

**DOI:** 10.1101/416255

**Authors:** Qin Qiao, Chakib Djediat, Hélène Huet, Charlotte Duval, Séverine Le Manach, Cécile Bernard, Marc Edery, Benjamin Marie

## Abstract

Among the diverse toxic components produced by cyanobacteria, microcystins (MCs) are one of the most toxic and notorious cyanotoxin groups. Besides their potent hepatotoxicity, MCs have been revealed to induce potential reproductive toxicity in various animal studies. However, little is still known regarding the distribution of MCs in the reproductive organ, which could directly affect reproductive cells. In order to respond to this question, an acute study was conducted in adult medaka fish (model animal) gavaged with 10 µg.g^-1^ body weight of pure MC-LR. The histological and immunohistochemical examinations reveal an intense distribution of MC-LR within hepatocytes along with a severe liver lesion in the toxin-treated female and male fish. Besides being accumulated in the hepatocytes, MC-LR was also found in the connective tissue of the ovary and the testis, as well as in oocytes and degenerative spermatocyte-like structures. Both liver and gonad play important roles in the reproductive process of oviparous vertebrates. This observation constitutes the first observation of the presence of MC-LR in the reproductive cell of a vertebrate model with *in vivo* study. Our results, which provide intracellular localization of MC-LR in the gonad, advance our understanding of the potential reproductive toxicity of MC-LR in fish.

## 1. Introduction

Recurrent cyanobacterial blooms frequently occur worldwide in eutrophic freshwaters. Various bloom-generating species of cyanobacteria can produce natural toxic components (cyanotoxins), and their blooms threaten human health as well as living organisms of the aquatic environment. Among cyanotoxins, microcystins (MCs) are the most prevalent cyanobacterial hepatotoxins, with 248 structural variants (Spoof and Catherine, 2017), being produced by at least six genera of cyanobacteria (Puddick et al., 2014). Among all these known variants, microcystin-LR (MC-LR) is considered to be the most common and potently toxic (Puddick et al., 2014).

MCs have difficulty to penetrate into vertebrate cells via passive transport due to their hydrophilic nature, and organic anion transporting polypeptides (OATPs) are known to be able to transport MCs through cell membranes (Campos and Vasconcelos, 2010). Once within the cell, MCs specifically inhibit eukaryotic serine/threonine protein phosphatases 1 and 2A, which causes the disruption of numerous cellular signals and processes (Fischer et al., 2005; MacKintosh et al., 1990). Although there have been 70 members of OATP superfamily identified in the database from humans, rodents and some additional species (Hagenbuch and Stieger, 2013), not all OATPs are capable to transport MCs through cell membranes. The capability of MC-transport has been demonstrated for OATP1A2 (mainly expressed in the vertebrate brain), liver-specific OATP1BA and OATP1B3 in humans (Fischer et al., 2005). In fish, liver-specific OATPs, such as OATP1d1 in little skate (Meier-Abt et al., 2007) and rtOATP1d1in rainbow trout (Steiner et al., 2014), have been reported to mediate MC-transport. Different types of OATP distributed in different tissue or organs possess varying levels of affinities and capacities for different MC variants (Fischer et al., 2010, 2005; Steiner et al., 2016). Liver is the organ that presents the highest tropism for MCs (so-called “target”), since it is rich in a few liver-specific OATP members possessing high affinities and capacities to MCs. Several studies have reported the acute hepatotoxicity of MCs characterized by hepatocellular apoptosis (Zhang et al., 2013), necrosis (Mattos et al., 2014), intrahepatic hemorrhaging (Hou et al., 2015), and cytoskeleton disruption (Zhou et al., 2015). Additionally, the distribution of MCs in the liver has been detected using immunohistochemical methods in mice (Guzman and Solter, 2002; Yoshida et al., 1998) and fish (Djediat et al., 2011; Fischer and Dietrich, 2000; Marie et al., 2012).

In addition to liver, MCs have been documented to distribute and accumulate in various fish organs including intestine, kidney, gill and gonad (Acuña et al., 2012; Djediat et al., 2010; Mezhoud et al., 2008; Trinchet et al., 2013, 2011). Among these organs, the gonad could be considered as a secondarily most important target of MCs, as several field studies reported also the presence of MCs in the gonad of fish. For instance, in Lake Pamvotis (Greece), the gonad of common carp (*Cyprinus caprio*) was reported to contain about 50 ng eq. MC-LR g^-1^ body weight (bw) using ELISA test (Papadimitriou et al., 2012), and in Lake Taihu (China), three variants of MC (MC-LR, -YR and -LR) were determined in the gonad of various fish species using a liquid chromatography-electrospray ionization-mass spectrum system (LC-ESI-MS). Particularly, silver carp (*Hypophthalmichthys molitrix*) and goldfish (*Carassius auratus*) were observed to contain high concentrations of MCs in the gonad (60 and 150 ng MCs g^-1^ DW, respectively) (Chen et al., 2009). In laboratories, the distribution of MC-LR in the gonad of medaka fish administered with pure toxin was also testified using a radiotracing method (Mezhoud et al., 2008). Although the accumulation of MCs in the gonad has been documented previously through different methods, little is known about the localization of MCs within the gonad tissue, or about the intracellular distribution of MCs. Only one field study reported the presence of MCs in the gonadal somatic tissue, but apparently not in the oocytes in common bream (*Abramis brama*) using immunohistochemical method (Trinchet et al., 2013). To date, it is still hard to know whether MC could enter the reproductive cells or only accumulate in the conjunctive tissue of gonad. Although not the most sensitive detection method, the immunohistochemical examination is a well-established way of providing the precise subcellular localization of MCs in the gonad, which could largely advance our current knowledge of the directly toxic effects of MCs on the gonad.

In the present study, in order to investigate subcellular distribution of MCs in fish gonads, an immunohistochemical assay with light microscopy and electron microscopy was conducted in adult medaka fish that were gavaged with acute doses of MC-LR (10 µg.g^-1^ bw of MC-LR, 1 h exposure). Localization of the toxin in the gonad was showed by MC-LR-specific antibodies, MC10E7, that specifically binds to the arginine at position 4 of MC-LR (Zeck et al., 2001). In addition, the toxin distribution and histopathological change in the liver (the first target of MCs) were also investigated in order to compare the toxic effect in liver and gonad.

## 2. Materials and methods

### 2.1 Chemical and reagent

MC-LR, purchased from Novakit^®^ (Nantes, France), was selected as the model MC for the present experiment. Five hundred µg of MC-LR was dissolved in 500 µL of ethanol and 500 µL of water. The ethanol was evaporated with a Speedvac. The concentration of MC-LR in the obtained solution was quantified using a commercial Adda-specific AD4G2 ELISA test (Abraxis), and then adjusted to be 1 µg.µL^-1^.

### 2.2 Fish maintenance, exposure and sampling

The experimental procedures were conducted in accordance with the European Union regulations and the supervision of the Cuvier’s ethical committee of the Museum National d’Histoire Naturelle (MNHN) concerning the protection of experimental animals. Five-month-old medaka fish (*Oryzias latipes*) of the inbred Cab strain maintained in the lab was used for this experiment. Female and male fish were maintained in 2 glass aquaria, respectively, filled with a mixture of tap water and reverse osmosis filtered water (1:2) in a flow-through system for aeration and filtration, in a temperature controlled room (25 ± 1 °C), with a 12 h:12 h light:dark cycle. Fish were fed three times a day with commercial dry bait for juvenile salmons.

Eight females and eight males were randomly selected from the aquaria. The fish were anesthetized with tricaine (150 mg.L^-1^) before gavage. Five females and five males were gavaged with 5 µL of MC-LR solution (1 µg.µL^-1^) per fish, representing about 10 µg.g^-1^ bw of MC-LR (average body weight is 0.52 g, the body weight of individual fish is shown in Supplementary table 1). This concentration is modified according to a few previous studies in which medaka fish exhibited noticeable tissue damage and toxin presence in liver upon exposure to 5 µg.g^-1^ bw of MC-LR (Djediat et al., 2011, 2010). Two females and two males were gavaged with 5 µL water as the non-toxin control, and 1 female and 1 male without any treatment were used as the non-gavage control. After 1 h oral exposure, fish were anesthetized with tricaine, sacrificed, and the liver and gonad were collected on ice. One tissue was cut into 2 parts, one part was immediately fixed with formaldehyde fixing solution (Supplemental material 1) according to the protocol provided by the histopathological platform of Ecole Nationale Vétérinaire d’Alfort (ENVA), and the other part was fixed with paraformaldehyde fixing solution (Supplemental material 2) following the instruction of the electron microscopy platform of Muséum National d’Histoire Naturelle (MNHN).

### 2.3 Histopathological observation and immunolocalization of microcystins

In the platform of ENVA, the liver and gonad samples fixed in formaldehyde fixing solution at 4 °C for 48 h were dehydrated in successive baths of ethanol (from 70 to 100%) and embedded in paraffin. Blocks were cut into 4 µm thick sections. The sections of each sample were deposited on 3 slides. One slide was stained with hematoxylin-eosin-saffron (HES) staining for histopathological observation. Another slide (only liver sample) was stained with periodic acid-Schiff (PAS)/Alcian blue staining for the assessment of hepatic glycogen reserve. The glycogen reserve analysis performed by ImageJ software (version 1.51d) estimated the hepatic glycogen quantity as a surface percentage of the purple-red pixels on 5 randomly selected photos (200 × magnification, photo size 1388×1040 pixels, as seen in Figure 1) per individual. The last slide, for immunolocalization of MC, was incubated with a monoclonal antibody to MC-LR (MC10E7, Alexis) that recognizes all MCs with Arg in position 4 (dilution 1:4000). The immunolabeling was routinely performed with an automated Module Discovery XT (Ventana, Tuscon, USA) using a colorimetric peroxidase-specific staining with diaminobenzidine (DAB), a substrate that produces a highly insoluble brown precipitate. The slides were counterstained with hematoxylin in the end. The secondary antibody control sections were prepared by skipping the first antibody MC10E7 incubation step. One MC-contaminated medaka liver sample from an acute high dose exposure in the lab previously was included as the technically positive control.

**Figure 1.**
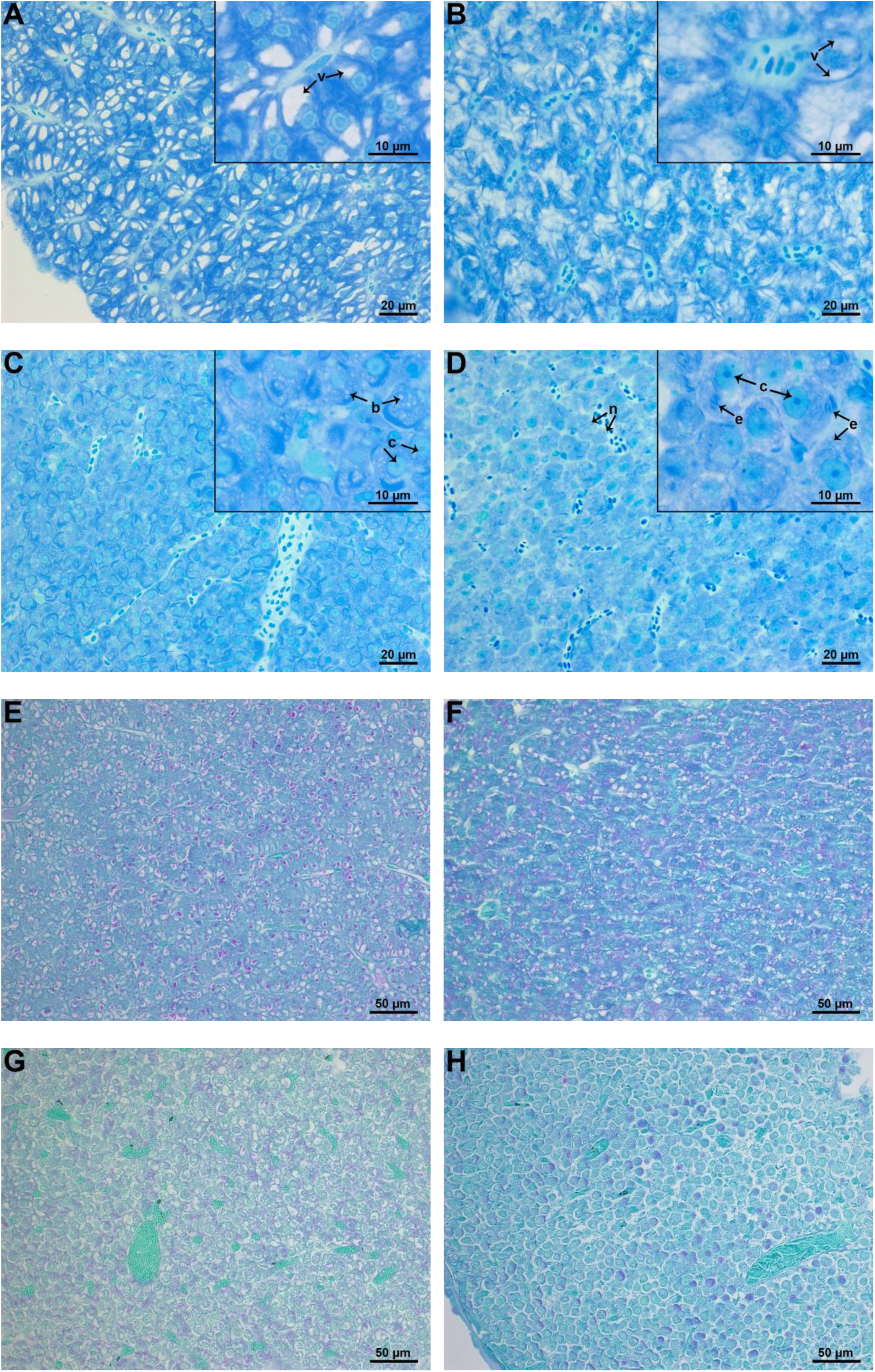
Representative photos of histopathological observation of medaka liver under a light microscope. A-D: resin sections (0.5 µm thick) of medaka liver with toluidine blue staining. Female control (A): abundant and distinct distribution of reserve vesicles (v); Females exposed to 10 µg.g^-1^ bw of MC-LR (C): disintegration of the cord-like parenchymal organization, rounded hepatocytes containing various small bubbles (b), loss of reserve vesicles, nuclear membrane breakdown, chromatin concentration (c); Male control (B): abundant and distinct distribution of reserve vesicles (v); Males exposed to 10 µg.g^-1^ bw of MC-LR (D): disintegration of the cord-like parenchymal organization, swollen and rounded hepatocytes, nuclear chromatin concentration (c), and increase in endoplasmic reticulum (e) and loss of nuclei (n). E-H: paraffin sections (4 µm thick) of medaka liver with PAS staining. Female (E) and male (F) control: abundant and distinct distribution of glycogen-reserve vesicles (red-purple); Females (G) and males (H) exposed to 10 µg.g^-1^ bw of MC-LR: less glycogen-reserve vesicles compared with the control group.

In the platform of MNHN, the liver and gonad samples fixed in paraformaldehyde fixing solution at 4 °C overnight were dehydrated in successive baths of ethanol (from 30 to 100%) and embedded into the Unicryl resin. Blocks were cut into 0.5 µm-thick semi-thin sections (stained with toluidine blue staining) and 60 nm-thick ultrathin sections for histopathological observation and immunolocalization of MC with electron microscopy, respectively. The 60 nm-thick ultrathin sections were collected by gold grids. Then, the sections were incubated with the same primary antibodies (MC10E7) at 4 °C overnight and rinsed, then incubated with a secondary antibody coupled to gold nanoballs (6 or 10 nm of diameter) at room temperature for 1 h. After being well rinsed with distilled water, the sections were stained with a saturated solution of uranyl acetate in 50% ethanol and then observed under an H-7700 Hitachi (Tokyo, Japan) transmission electron microscope.

## 3. Results

During the 1 hour of post-gavage, all MC-treated fish exhibit abnormal swimming behaviors, such as difficulty in moving or losing body balance. In the contrary, the fish of control group all swim normally. Histopathological and immunohistochemical observation results of each individual are summarized in Table 1, which shows visible toxic effects and clear toxin distribution in nearly every MC-treated fish, but not in any control fish.

**Table 1.**
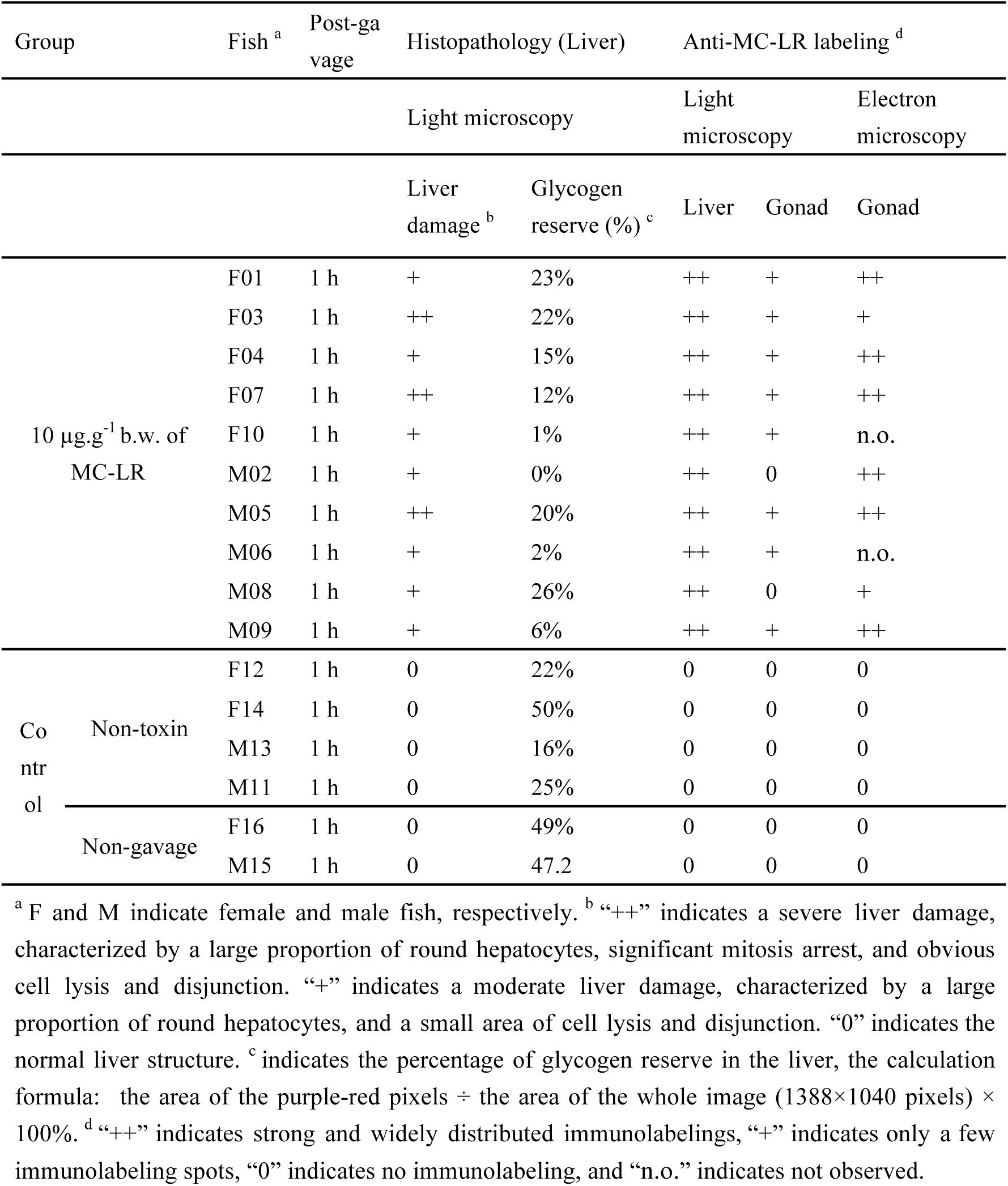
Histopathological and immunohistochemical observation of each individual liver and gonad

### 3.1 Histopathological effects with light microscopy

In the liver of all control fish, hepatocytes present compact and characteristic cord-like parenchymal organization and a distinctive central nucleus with a prominent nucleolus (Figure 1 A and B). The cytoplasm of hepatocytes contains mainly glycoprotein and/or glycogen stores that are intensively stained in red-purple with PAS staining (Figure 1 E and F). In contrast, the livers of MC-LR treated fish exhibit noticeable tissue lesions characterized by a disintegration of the parenchymal organization and significant cell lysis. The hepatocytes all lose their polyhedral shape and become rounded and swollen. Within the hepatocytes, the reserve vesicles disappear, the cytoplasm presents various small bubble-like structures, and the nuclear chromatin becomes dense. Some hepatocytes exhibit a breakdown of the nuclear membrane, mitosis arrest, and dilation of endoplasmic reticulum (Figure 1 C and D). In addition, the hepatic intracellular glycoprotein and/or glycogen quantity seems to decrease, and the mean values of glycogen percentage for female controls, male controls, female toxin-treated fish and male toxin-treated fish are 15%, 11%, 40% and 29%.

For the gonad, there is no apparent cellular difference between the toxin-treated fish and the control ones.

### 3.2 Immunolocalization of MC-LR in the liver and the gonad

Through light microscopy, all livers of MC-LR treated fish exhibit a strong positive signal of MC-LR specific antibodies (Figure 2 A) whereas the secondary antibody control does not display any positive signal (Figure 2 B), being similar to control fish in which no immunolabeling is observed (Figure 2 G). For the toxin treated fish, the brown immunolabelings are evenly distributed in hepatocytes, but not in erythrocytes which are only stained in blue by hematoxylin. Within hepatocytes, the immunolabeling of MCs is showed in the cytoplasm and more intense labeling is observed in nuclei (Figure 2 A). Immunogold electron microscopy also shows that the black nanoballs (immunolabelings) are localized in the cytoplasmic inclusion and the nucleus (Figure 3 C) of hepatocytes of MC-LR treated fish, being particularly intense in the lysis area (Figure 3 B). MC-LR is also distributed intensely in residual bodies of the macrophage (Figure 3 D), whereas no immunolabeling is observed in the control fish (Figure 3 A) or in the secondary antibody control sections.

**Figure 2.**
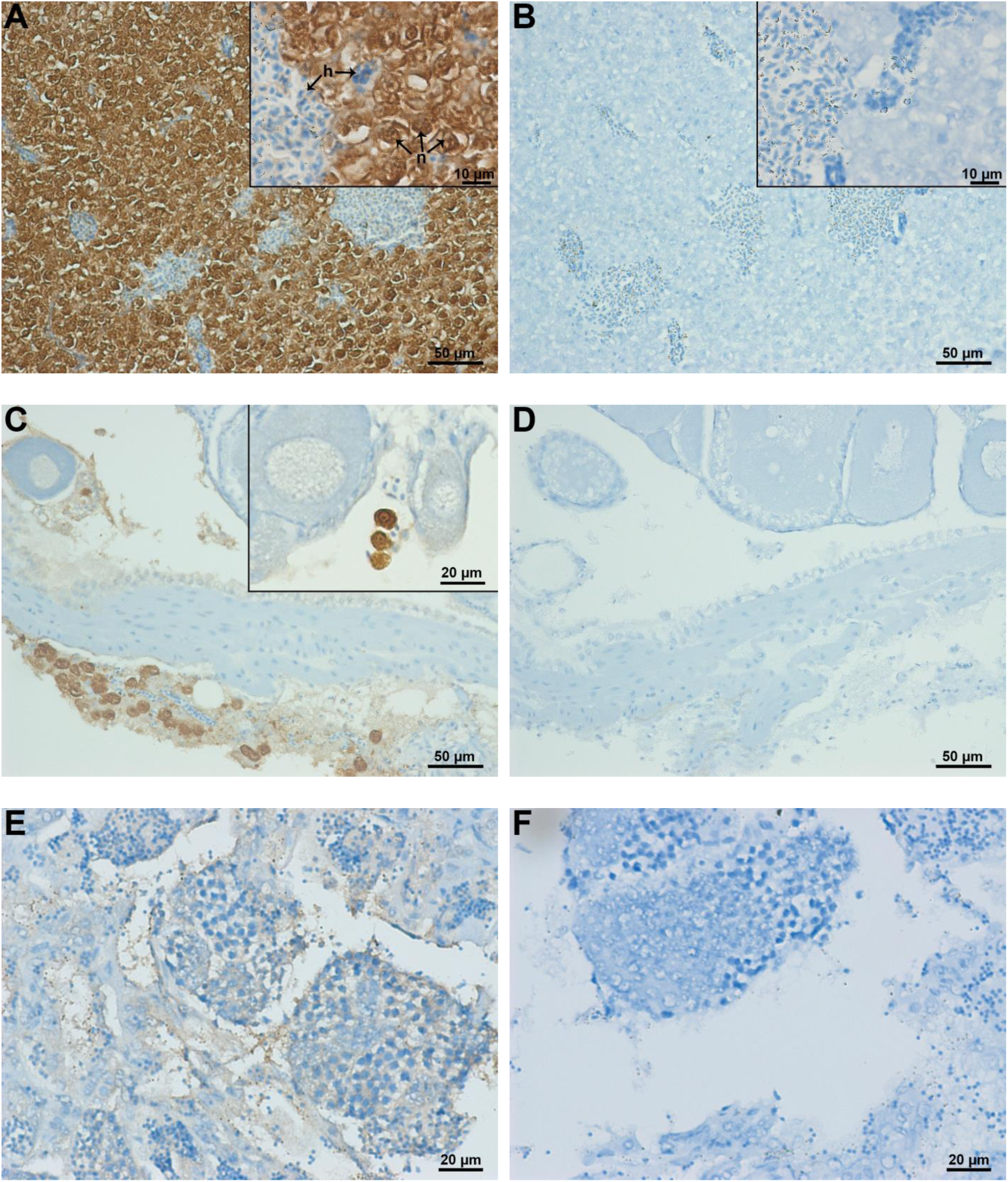
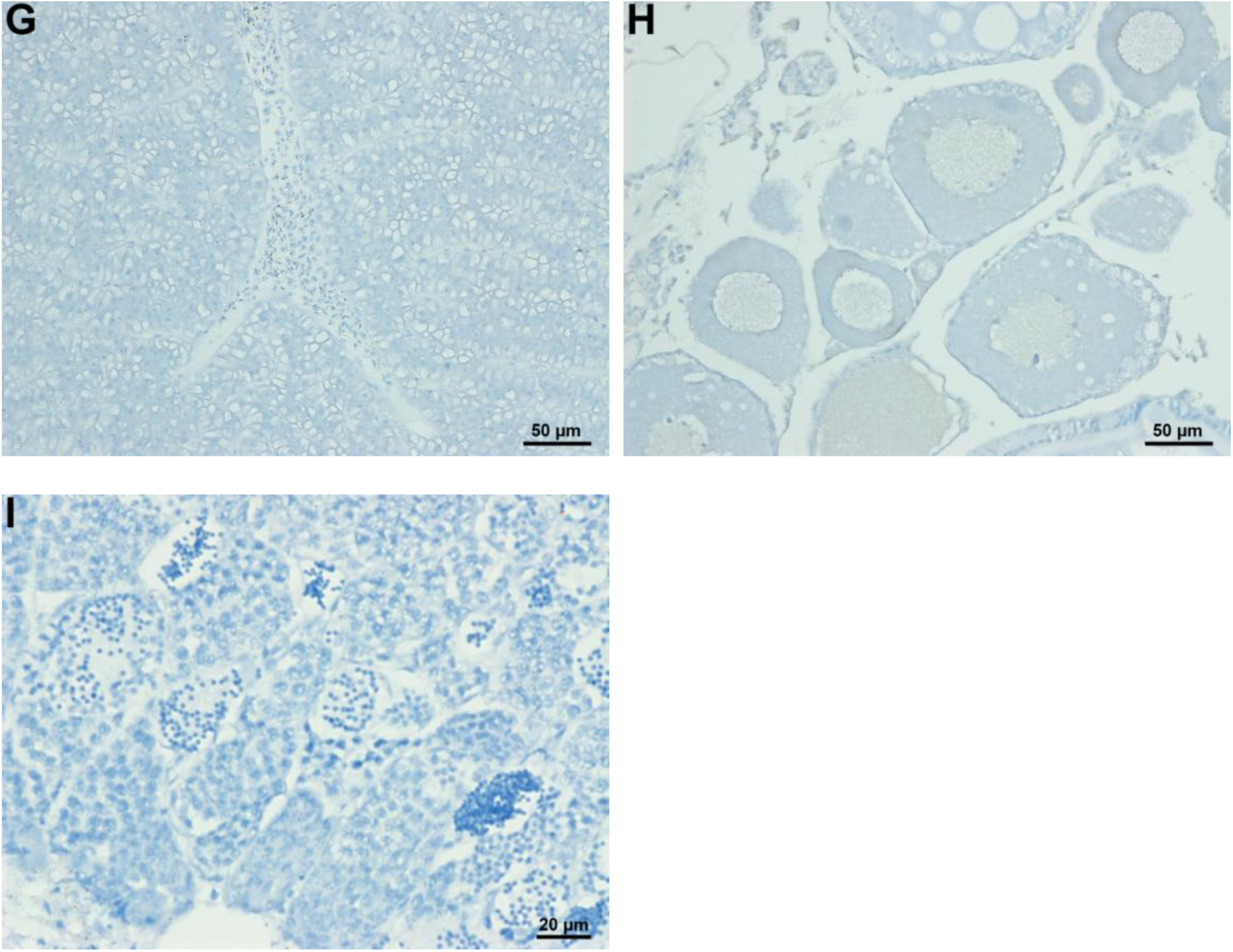
Representative photos of immunolocalization of MC-LR in the liver and gonad of toxin-treated fish through light microscopy. Sections of the toxin-treated female liver (A), ovary (C), and testis (E), stained with both toluidine blue and MC-LR immunolabeling (MC10E7), revealed with peroxidase specific reaction of DAB. n, nucleus; h, hematoxylin; The secondary antibody control sections of the toxin-treated female liver (B), ovary (D), and testis (F), stained with toluidine blue; Sections of non-toxin control female liver (G), ovary (H), and testis (I), stained with toluidine blue.

**Figure 3.**
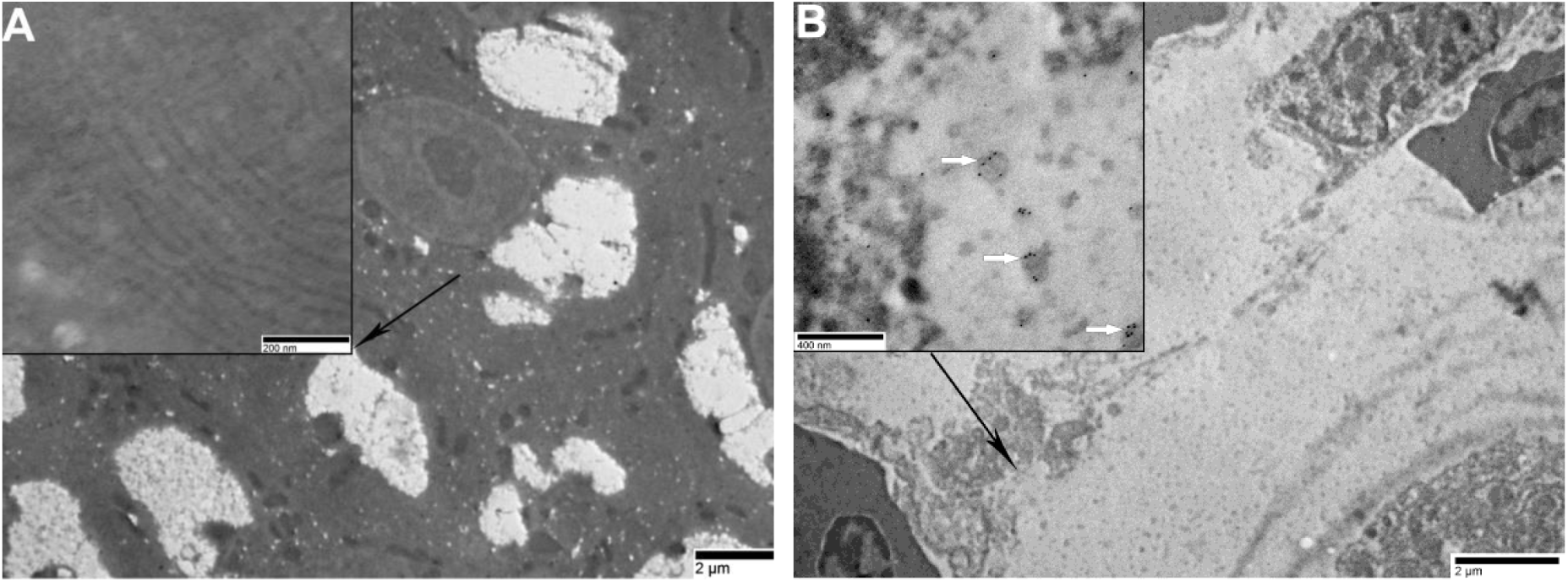
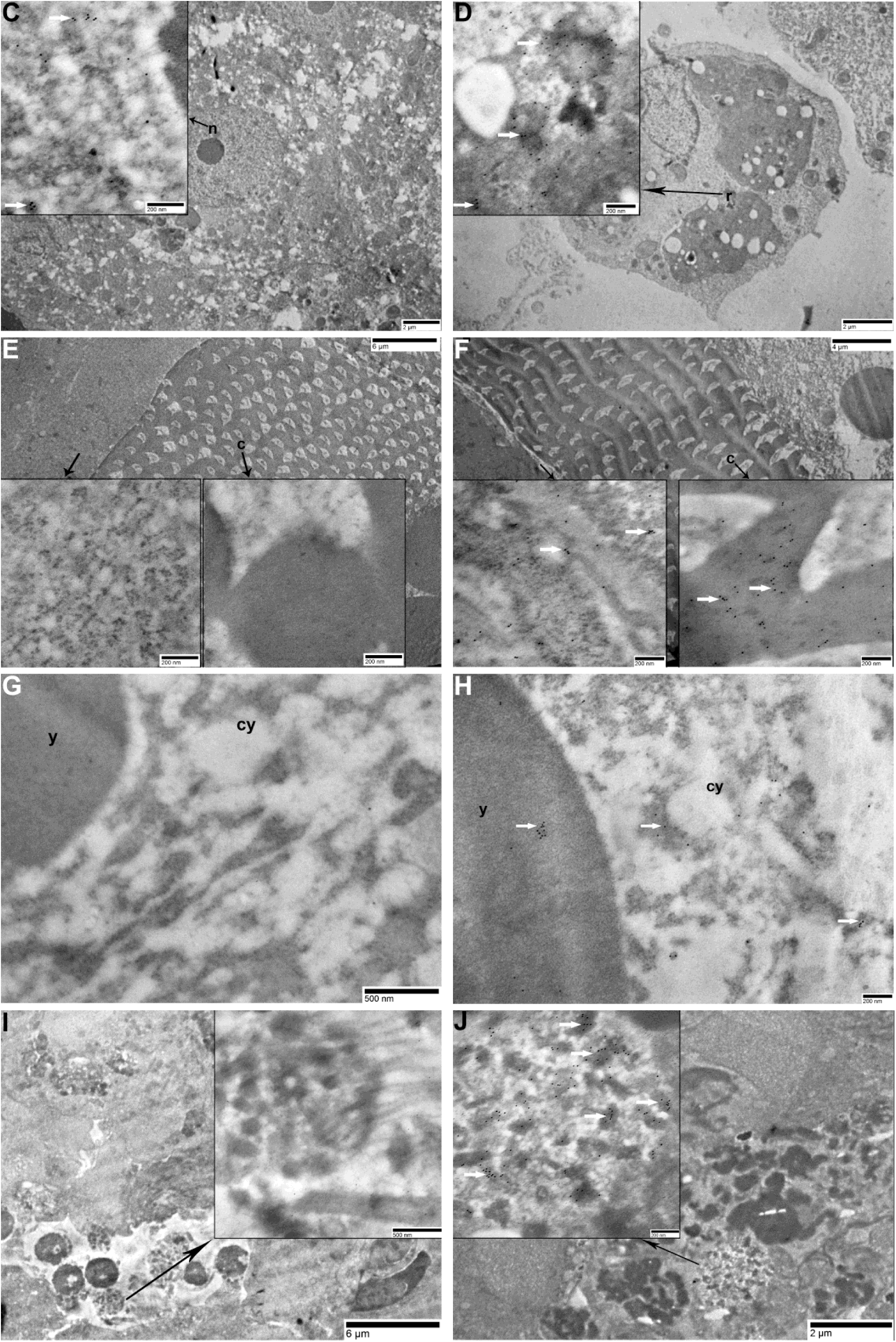
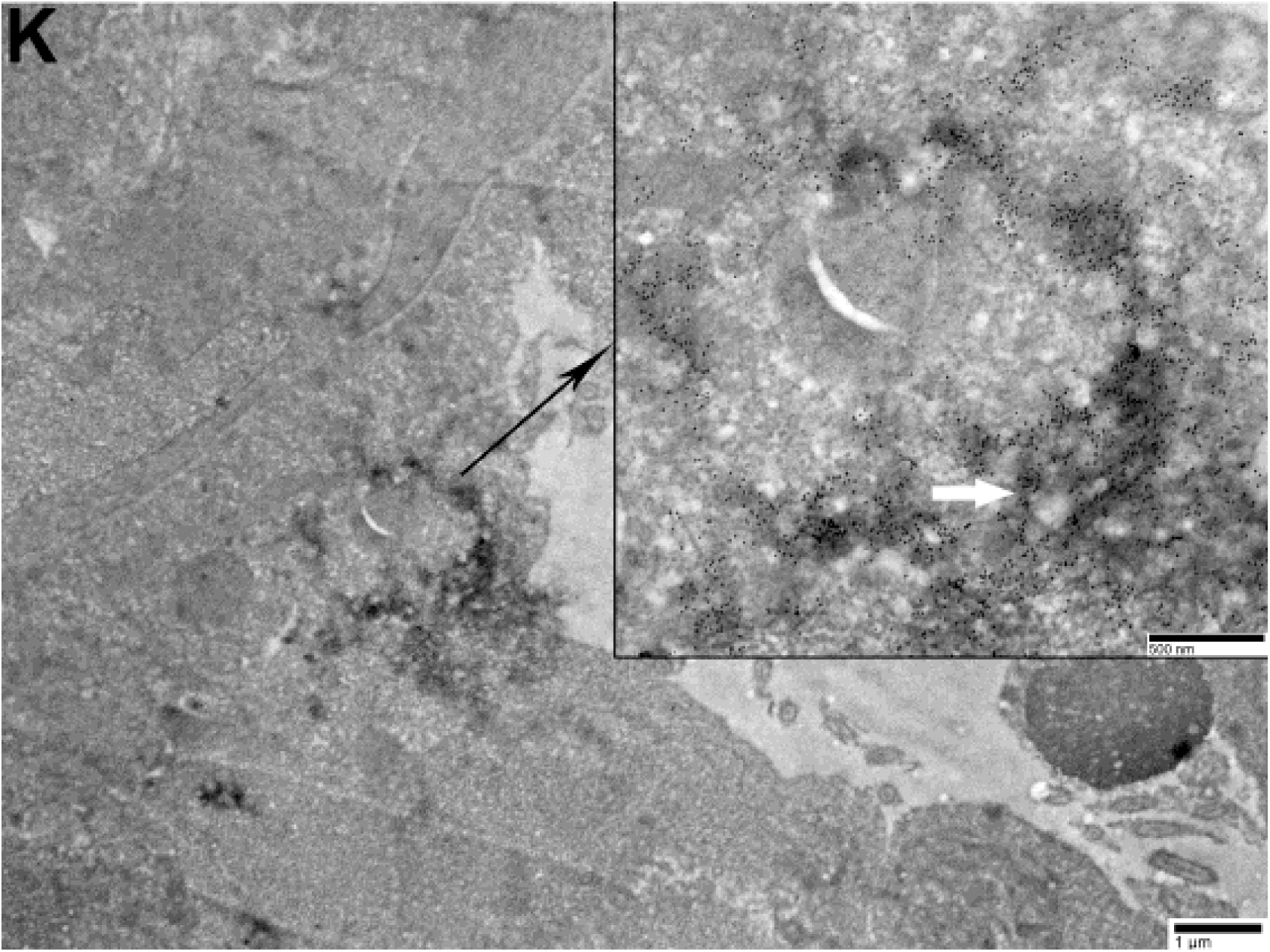
Representative photos of immunolocalization of MC-LR in the liver and gonad of toxin-treated fish through immunogold electron microscopy. The liver (A), ovary (E and G) and testis (I) of control fish: no clear immunolabeling. The liver (B-D), ovary (F-H) and testis (J and K) of the fish exposed to 10 µg.g^-1^ bw of MC-LR, showing the immunolabeling of MC-LR indicated by the white arrow. For liver, immunolabelings of MC-LR are clearly observed in the lytic residual (B) and nuclear (C) of hepatocyte and the macrophage (D). For ovary, the immunolabeling is found in the connective tissue (left) and the chorion (c) of the oocyte (right), as has been shown in F, and less labelings are observed in the yolk vesicle (y) and the cytoplasm (cy) of the oocyte (H). For testis, the labeling is intensively distributed in some unidentified round structures, which appeared to be degenerated germ cells (J) and the connective tissue of some certain area (K)

By light microscopy observation of the ovary of toxin-treated fish, clear immunolabelings are detected in the connective tissue and some round cells with above 10 µm in diameter, being probably the early stage of oocyte (Figure 2 C), whereas the secondary antibody control does not display any positive signal (Figure 2 D), being similar to the control fish in which no immunolabeling is observed (Figure 2 H). Moreover, electron microscopy shows that the immunogold labelings are not only present in the gonadal somatic cells, but also distributed intensively in the chorion of oocytes (Figure 3 F), being less intense in the yolk vesicles and cytoplasm of oocytes (Figure 3 H), whereas no immunolabeling is observed in either the control fish (Figure 3 E and G) or the secondary antibody control sections.

On light microscope, weaker immunolabelings are found in the testicular connective tissue of a few toxin-treated male fish (Figure 2 E), whereas the secondary antibody control does not display any positive signal (Figure 2 F), being similar to the control fish in which no immunolabeling is observed (Figure 2 I). Immunogold electron microscopy shows a remarkable labeling in some round unidentified structures, being probably some degenerated germ cells (Figure 3 J), as well as in the connective tissue of certain area nearby seminiferous tubules (Figure 3 K), whereas no immunolabeling is observed in either the control fish (Figure 3 I) or the secondary antibody control section.

## 4. Discussion

MC-treated female and male fish in this study show the characteristic hepatocellular changes and the depletion of glycogen reserve that have been observed previously in fish (Djediat et al., 2011; Fischer et al., 2000; Fischer and Dietrich, 2000; Marie et al., 2012; Mezhoud et al., 2008) and mice (Guzman and Solter, 2002; Yoshida et al., 2001, 1998) acutely administered with MCs or MC-containing cyanobacteria extracts. In fact, the observed liver damage in the present study is also similar to some of those described in the fish after chronic intoxication with MCs (Acuña et al., 2012; Qiao et al., 2016; Trinchet et al., 2011). Trinchet and her colleagues observed an increase in rough endoplasmic reticulum in hepatocytes of medaka fish chronically exposed to MC-LR (5 µg.L^-1^, 30 d), which is also found in the male toxin-treated medaka in the present study. This could be associated with the induction of detoxification process. Another chronic in-house study reported a noticeable hepatic glycogen store depletion in medaka fish upon exposure to MC-LR (5 µg.L^-1^, 28 d) (Qiao et al., 2016), which is consistent with the acute intoxication effect of MC-LR observed here, and the increased energy needs for the MC detoxification process might be the cause.

Severe liver damages are often accompanied by remarkable accumulation of MCs in the liver. The present study shows an intense distribution of MC-LR in the lytic hepatocyte of the toxin-treated medaka fish through the immunohistochemistry. MCs have been reported to mostly localize in the cytoplasm of hepatocytes where they disturb a sequence of phosphorylation/dephosphorylation-dependent biochemical reactions, resulting in the disruption of multiple cellular processes, and consequently causing the deformation of hepatocytes, apoptosis and lysis (Campos and Vasconcelos, 2010). Furthermore, a high concentration of MC-LR is also observed in the hepatic nucleus here, which is consistent with the result provided in a few previous acute studies (Fischer et al., 2000; Guzman and Solter, 2002; Yoshida et al., 1998). Protein phosphatases PP1 and PP2A, one of the main molecular “target” of MCs, localize in both nucleus and cytoplasm (Shenolikar, 1994). Therefore, the observed nuclear accumulation of MC-LR could be the consequence of the MC specific antibodies binding to the MC-LR-PP1/PP2A adducts localized in the nucleus.

In the present study, MC-LR is also observed to be present in the gonad of MC-treated fish. For females, MC-LR is localized in the connective tissue of ovary, as well as in some maturing oocytes. The presence of MC in the connective tissue of fish ovary has been reported previously in common bream collected from MC-contaminated lakes (Trinchet et al., 2013). It seems that MCs are transported through the bloodstream into gonads, in which they can firstly accumulate in the connective tissue, and then transported into various gonadal somatic cells, even reproductive cells. Our result indeed exhibits an apparent distribution of MC-LR in the oocyte of vertebrates with *in vivo* condition for the first time, and the toxin is clearly localized in the yolk and the cytoplasm of the oocyte, being particularly highly intensive in the chorion. Through the immunochemistry, MC was once reported to be present in the oocyte of snail upon exposure to MC-LR for 5 weeks (Lance et al., 2010). But it is unclear how MC-LR enters into oocytes since the oocyte plasma membrane seems to lack the OATPs possessing MC-transport capability. It can be assumed that MCs may be transported into oocytes as protein-bound forms along with protein import during oogenesis, since the exchange of a big amount of protein and other materials are occurring prior to complete chorion maturation, or even simply through the passive diffusion of a small quantity.

For MC-treated male fish, immunogold electron microscopy reveals an intense distribution of MC-LR in the connective tissue of the testis in some certain area, as well as in some degenerated spermatocyte-like structure. It is uncertain whether the observed degenerated structure is caused by the toxic effect of MC-LR, or is associated with the natural apoptotic loss, since a certain level of apoptosis or degeneration in germ cells is normal in fish testis (Schulz et al., 2010). The observed high accumulation of MCs in a certain area of connective tissue still implies a potential germ cell damage induced by MC exposure. The connective tissue of the testis consists of massive fibers and various types of gonadal somatic cells, such as Leydig cells and Sertoli cells. They are important functional cells in the testis, being responsible for synthesizing and secreting androgens that are essential for the development and maturation of germ cells or serving as channels for the transport of nutrients into the growing germ cells (Dietrich and Krieger, 2009). Therefore, the high accumulation of MC-LR in the connective tissue and its subsequent toxic effect on gonadal somatic cells may indirectly disturb normal spermatogenesis process, affecting the normal reproductive process. In the present study, we do not observe any histopathological modification in the spermatocyte of MC-treated fish, which seems to be due to the short exposure time (1h). In a previous in-house study, MC-LR has been reported to result in the disturbance of spermatogenesis in medaka fish following a chronic exposure (Trinchet et al., 2011). Considerable testicular injury and spermatocyte abnormality have been found in several species of vertebrate model animals, such as mouse, rat and fish, upon exposure to MCs acutely or chronically (Li et al., 2008; Su et al., 2016; X. Wang et al., 2013; Zhao et al., 2012; Zhou et al., 2013). However, to date, according to *in vivo* studies using these vertebrate models, no evidence regarding the incorporation of MCs into spermatocytes has been reported yet. Only one study showed a clear distribution of MC-LR on the tubal wall of seminiferous tubules of the MC-treated rats through the immunofluorescence method (injection with 300 µg.kg^-1^ bw of MC-L for 6 d) (L. Wang et al., 2013). In the present study, MC-LR is not found in any clear spermatocytes of the toxin-treated fish through the immunogold labeling technique, which is consistent with the common knowledge of the organ distribution of the MC-transporting OATPs. The identified OATPs that possess strong MC-transport capabilities are highly abundant in the liver, but hardly known to be present in the cell membrane of reproductive cells. However, one *in vitro* study reported that MC-LR was able to immigrate into isolated rat spermatogonia (Zhou et al., 2012). Furthermore, in this study, at least 5 OATPs were detected at the mRNA level in spermatogonia and their expression level was affected by MC-LR, implying that these OATPs may involve in MC-transport into the reproductive cells. Therefore, it is noteworthy that some unidentified OATPs which possess MC-transport capabilities might be expressed in the reproductive cells at a relatively low level. But the information regarding the unidentified MC-transporting OATPs and their expression level in reproductive cells still remains very limited.

The present acute study indeed shows a clear subcellular distribution of MC-LR in the gonad of the toxin-treated fish. However, in another in-house chronic study, MCs were not detected in the ovary or testis of medaka fish following a balneation exposure to 5 µg.L^-1^ of MC-LR for 30 days by using the same immunolocalization techniques (Trinchet et al., 2011). There are at least two causes that may account for this discrepancy. Toxin concentration, detoxification process and exposure time are crucial for such immunolabeling results. Ten µg.g^-1^ bw of MC-LR used in the present study is a quite high concentration (the highest LD_50_ value of MCs by i.p. injection is about 1.5 µg.g^-1^ bw in fish) (Malbrouck and Kestemont, 2006), and the short time of exposure (1 h) largely reduced the possible toxin excretion process through liver detoxification, together leading to a sufficient quantity of toxin that could access to gonad through blood stream and be detected by the immunohistological method. Besides, it is worth to mention that the process of histological section preparation may affect the result, since a fraction of MC-LR (mainly free MC-LR) was removed during the dehydrating process of the section preparation. The immunostaining here indeed detects MC-LR-PP1/PP2A adducts mostly, due to the strong affinity of MC-LR to protein phosphatases PP1/PP2A (Yoshida et al., 2001). Dmet-Asp and D-Glu residues of MC-LR play important roles in forming the MC-LR-PP1/PP2A adducts (Campos and Vasconcelos, 2010).

## 5. Conclusion

Our histological and immunohistochemical results reveal that both liver and gonad are significantly affected by MC-LR exposure. An intense distribution of MC-LR within hepatocytes along with a severe liver damage attests to the potent hepatotoxicity of MC. The immunohistochemical results show that, besides being accumulated in the hepatocytes, MC-LR is also found in the connective tissue of the gonad, as well as in the reproductive cell (oocytes). This finding constitutes the first observation of the presence of MC in the reproductive cell in vertebrate model animals with *in vivo* condition. Both liver and gonad play important roles in the reproductive process of oviparous vertebrates. Our results of the present acute study, which provide a distinct subcellular localization of MC-LR in the liver and gonad, contributes to a better understanding of the potential reproductive toxicity of MC-LR at the histopathological level, favoring the characterization of underlying mechanisms. Meanwhile, the penetration of MCs into the reproductive cell suggests a possible transferring of MCs from adults into offspring which could cause a big issue for the population of aquatic organisms. The further investigation concerning this perspective is needed to advance our current knowledge of the protection of aquatic organism populations, as well as human beings from the widespread MCs in the freshwater body.

## Acknowledgments

This work was supported by grants from the CNRS Défi ENVIROMICS “Toxcyfish” project and from the ATM “Cycles biologiques: evolution et adaptation” of the MNHN to Dr. Benjamin Marie. Qin Qiao PhD is founded by the China Scholarship Council. We thank the Amagen platform for providing medaka fish Cab strain, the microscopy platform of ENVA and MNHN for the histopathology and immunolocalization techniques. We also thank Marie-Claude Mercier for its administrative support.

## Author Contribution

Q.Q., C.D., H.H., C.B., M.E. and B.M. conceived the experiments, Q.Q., C.D., H.H, C.D., S.L.M. and B.M. conducted the experiments, Q.Q., C.D., H.H and B.M. analyzed the results. All authors reviewed the manuscript.

The authors declare no competing financial interest.

## List of Supplementary materials

**Supplementary material 1.** Formaldehyde fixing solution (100 mL) provided by the platform of ENVA

|  |  |
| --- | --- |
| Formaldehyde (37-40%) | 20 mL |
| Glycerol | 10 mL |
| Glacial acetic acid | 10 mL |
| Absolute ethanol | 30 mL |
| Distilled water | 30 mL |

**Supplementary material 2.**
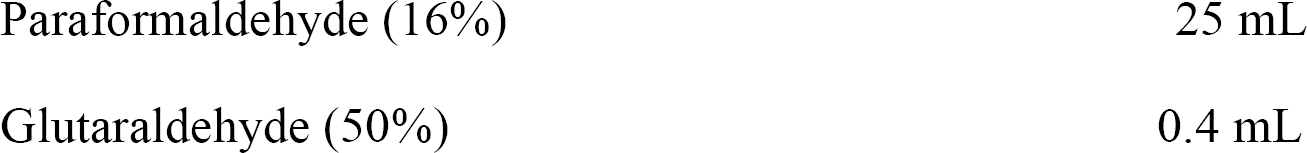
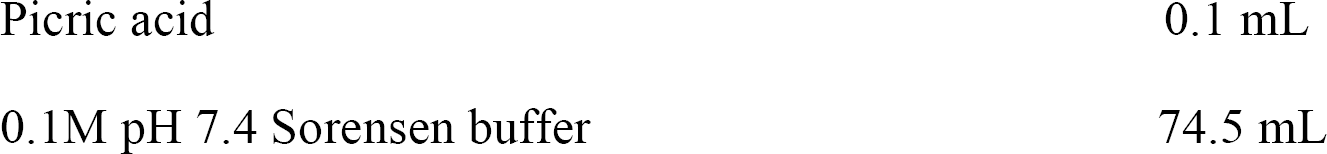
Paraformaldehyde fixing solution (100 mL) provided by the platform of MNHN

**Supplementary Table 1.**
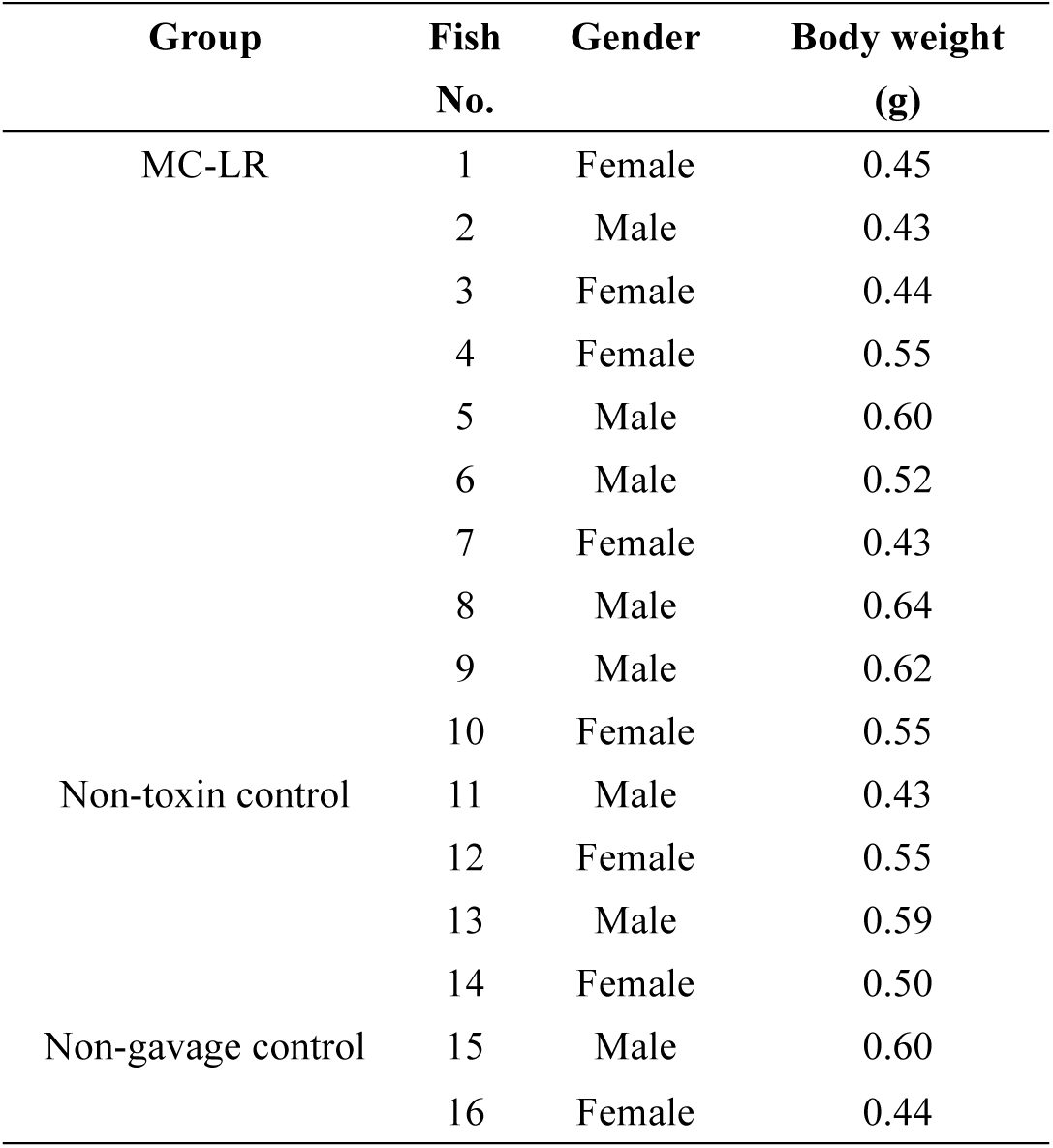
The body weight of individual fish in different treatment groups

| Group | Fish No. | Gender | Body weight (g) |
| --- | --- | --- | --- |
| MC-LR | 1 | Female | 0.45 |
|  | 2 | Male | 0.43 |
|  | 3 | Female | 0.44 |
|  | 4 | Female | 0.55 |
|  | 5 | Male | 0.60 |
|  | 6 | Male | 0.52 |
|  | 7 | Female | 0.43 |
|  | 8 | Male | 0.64 |
|  | 9 | Male | 0.62 |
|  | 10 | Female | 0.55 |
| Non-toxin control | 11 | Male | 0.43 |
|  | 12 | Female | 0.55 |
|  | 13 | Male | 0.59 |
|  | 14 | Female | 0.50 |
| Non-gavage control | 15 | Male | 0.60 |
|  | 16 | Female | 0.44 |

